# Ultrastructural expansion microscopy (U-ExM) visualization of malaria parasite dense granules using RESA as a representative marker protein

**DOI:** 10.1101/2024.11.06.622397

**Authors:** Junpei Fukumoto, Takafumi Tsuboi, Eizo Takashima

## Abstract

Dense granules (DG) are understudied apical organelles in merozoites, the malaria parasite stage that invades erythrocytes. Only six proteins have been identified which localize to DGs, despite that DG proteins play crucial roles in multiple steps of intraerythrocytic parasite development. To develop a tool for investigating DG structure and function, this study applied ultrastructural expansion microscopy (U-ExM) to visualize the ring-infected erythrocyte surface antigen (RESA) in *Plasmodium falciparum* merozoites. Merozoites were expanded to approximately four times their original size, allowing the identification of DGs without the need for electron microscopy. RESA localization in merozoite DGs was confirmed by staining with a combination of anti-RESA mAb and protein staining by NHS-ester. The translocation of RESA to the infected erythrocyte membrane was also observed in early ring-stage parasites. These results are in good agreement with the RESA localization reported using immunoelectron microscopy (IEM). By using U-ExM, the identification of novel DG proteins will be facilitated without time-consuming IEM, thereby contributing to describing erythrocyte parasitism by *P. falciparum*.

---

Human malaria is caused by *Plasmodium* spp. protozoa, which are transmitted by *Anopheles* mosquitoes. Malaria pathogenesis involves a complex interaction of parasite proteins that facilitate invasion, growth, and subsequent egress from human erythrocytes. Merozoite apical organelles, including rhoptries, micronemes, and dense granules (DG), are of interest due to their critical roles in merozoite invasion and growth [1]. A comprehensive understanding of the functions of organellar proteins is pivotal for elucidating malaria parasite biology, pathogenesis, and protective immunity, as well as aiding the development of effective intervention strategies [1, 2]. However, the small size of merozoite apical organelles poses challenges for visualization using conventional techniques, such as confocal laser-scan microscopy. Therefore, immunoelectron microscopy (IEM) is widely used to investigate the subcellular localization of parasite molecules in merozoites. IEM itself entails challenges, such as the generation of quality antibodies, and the harsh processes for IEM sample preparation, including the fixation, dehydration, and embedding steps, which may disrupt target epitopes and, consequently, lead to the loss of reactivity of the antibodies used for target molecule staining.

To address the above limitations, recent studies have utilized ultrastructural expansion microscopy (U-ExM) to achieve high-resolution visualization of subcellular organelles and structures in asexual *Plasmodium falciparum* parasites – for example, microtubule, centriolar plaque, basal complex, inner membrane complex, mitochondrion, apicoplast, cytostome, rhoptry, microneme, endoplasmic reticulum, and Golgi apparatus [3]. However, DG in merozoites have not been characterized using U-ExM [4].

To date, only six *P. falciparum* merozoite DG proteins have been identified and characterized by IEM; namely, exported protein 2 (EXP2) [5], *Plasmodium* translocon of EXported protein (PTEX) 150 [5], exported protein 1 (EXP1) [6], parasitophorous vacuolar protein 1 (PV1) [7], liver-stage antigen 3 (LSA3) [8], and ring-infected erythrocyte surface antigen (RESA) [9]. Of these, we have used IEM to characterize EXP1, PV1, and LSA3. In addition, subtilisin-like serine protease (SUB1) was localized to a DG-like organelle termed the exoneme [10]. EXP2, PTEX150, and EXP1 form PTEX components that play essential roles in the trafficking of parasite proteins to the surface of infected RBCs. PV1 has a chaperone activity, and it has been suggested to help recruit cargo proteins in the parasitophorous vacuole (PV) to PTEX [11]. SUB1 is an essential protease for merozoite egress [10]. Although the function of LSA3 in the erythrocyte stage remains unknown, antibodies against LSA3 could inhibit merozoite invasion, suggesting its involvement in the invasion process.

DG and exoneme proteins are essential for biological processes in asexual-stage parasites. While these proteins could be potential drug and vaccine targets, they remain relatively understudied compared to rhoptry and microneme proteins. One of the technical obstacles to investigations is that these organelles are too small to identify once proteins are localized. Therefore, in this study, as a proof-of-concept of using U-ExM technology, we sought to visualize a well-known DG protein, RESA.

Ultrastructural expansion microscopy (U-ExM) allows fixed cells to be expanded roughly four times in size, thus allowing higher localization resolutions than with super-resolution microscopy. After fixation of *P. falciparum-*infected erythrocytes, samples were embedded in poly-acrylamide and acylate hydrogel, and then expanded isotopically with deionized water [3]. The samples were then stained with antibodies to visualize the target antigens, and amine-reactive NHS-ester conjugated with Alexa Fluor 405 was used to visualize protein density [12].

First, we confirmed a published result of U-ExM with IFA using rabbit polyclonal antibodies against a rhoptry body marker, rhoptry-associated protein 1 (RAP1), and a microneme marker, apical membrane antigen 1 (AMA1), combined with nuclear staining using SYTOX (Figure 1). RAP1 signals colocalized with club-shaped rhoptries, as reported [13]; and visualization was aided by the clear staining of the dense rhoptry structures with NHS-ester, obviating the need for rhoptry marker antibodies. In contrast, AMA1 co-localized to cloud-like NHS signals adjacent to the rhoptries.

**Fig 1.**
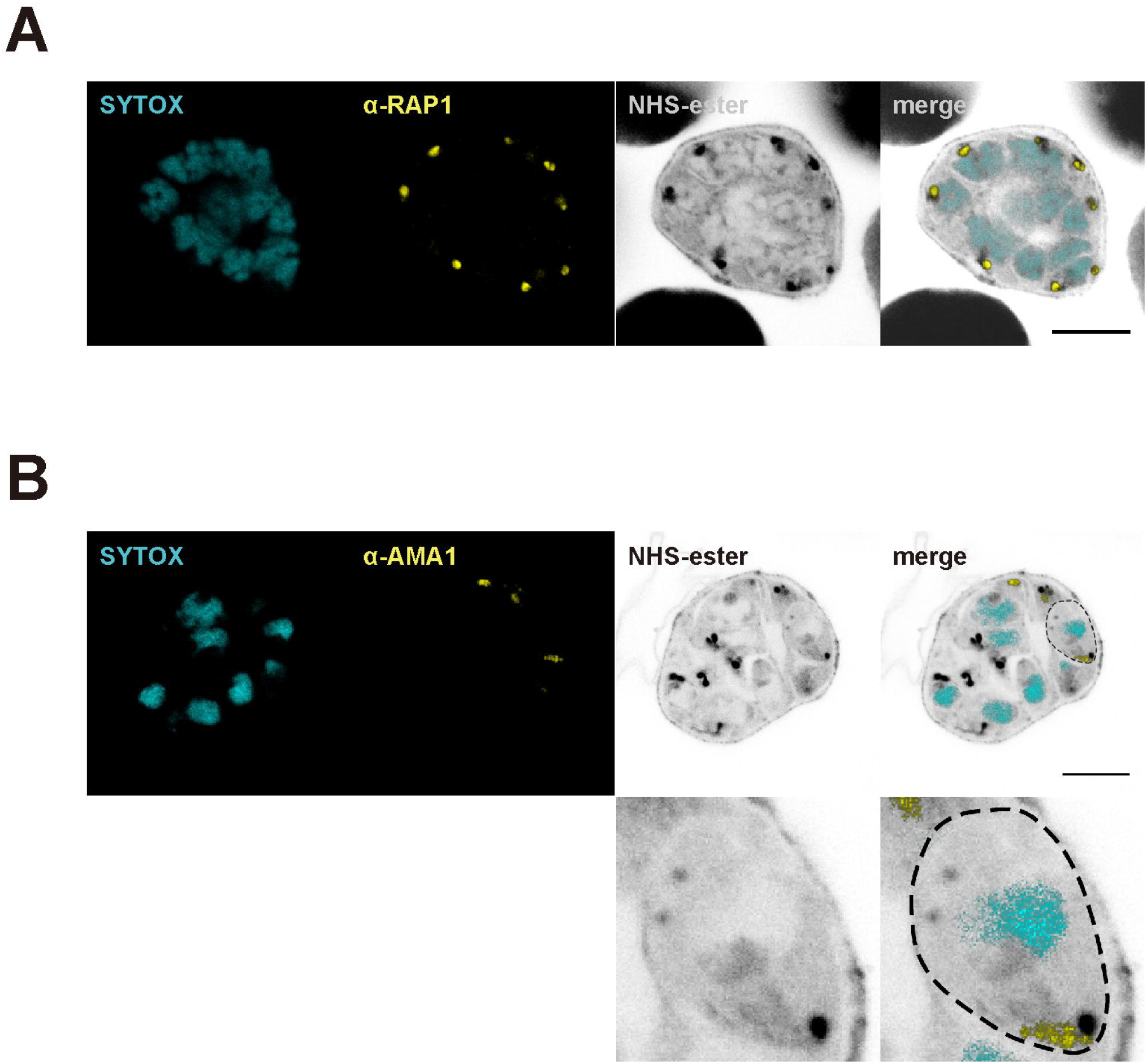
Anti-RAP1 and anti-AMA1 antibodies identified rhoptry and microneme organelles in expanded malaria parasites. Representative U-ExM images of parasites labeled with anti-RAP1 (A) or anti-AMA1 (B) antibodies. Protein amines and nuclei were visualized with NHS ester Alexa Fluor 405 and SYTOX Deep Red, respectively. RAP1 and AMA1 antibody labeling are shown in yellow. Protein density and nuclei are shown in grayscale and cyan, respectively. Scale bars = 10 μm.

Next, we attempted to observe DGs following immunolocalization with anti-RESA mAb. To confirm the anti-RESA mAb reactivity, we first conducted a conventional confocal laser-scan microscopy with the anti-RESA mAb. As previously shown, RESA localized on the infected erythrocyte cell membrane in the ring stage (Supplementary Figure 1, upper panels). Although RESA was observed in merozoites (Supplementary Figure 1, lower panels), one could not characterize it as a DG protein solely using conventional IFA. By U-ExM, RESA signal was found in merozoites in segmented schizonts, co-localized with protein-dense and spherical organelles distinct from the rhoptry or microneme upon NHS staining (Fig 2A), as observed by EM [3]. Some DGs were not stained with the anti-RESA mAb, suggesting that they were exonemes or a subpopulation of DGs where RESA is not localized [7]. In free merozoites, DGs with RESA signals were localized within most of the protein-dense structures distributed in the cytosol from the apical to the basal end (Fig 2B). In the ring stage, RESA is known to be secreted from DGs after invasion; and we observed it localized on the parasitophorous vacuole membrane (PVM) of ameboid ring stage parasites, accumulated at the tips of pseudopod-like structures, and faint staining on the infected erythrocyte membrane (Fig 2C, upper panels). In the immature schizont stage, RESA was localized on the PVM; however, infected erythrocyte membrane staining disappeared (Fig 2C, lower panels). These results are concordant with previous IEM studies [14, 15].

**Fig 2.**
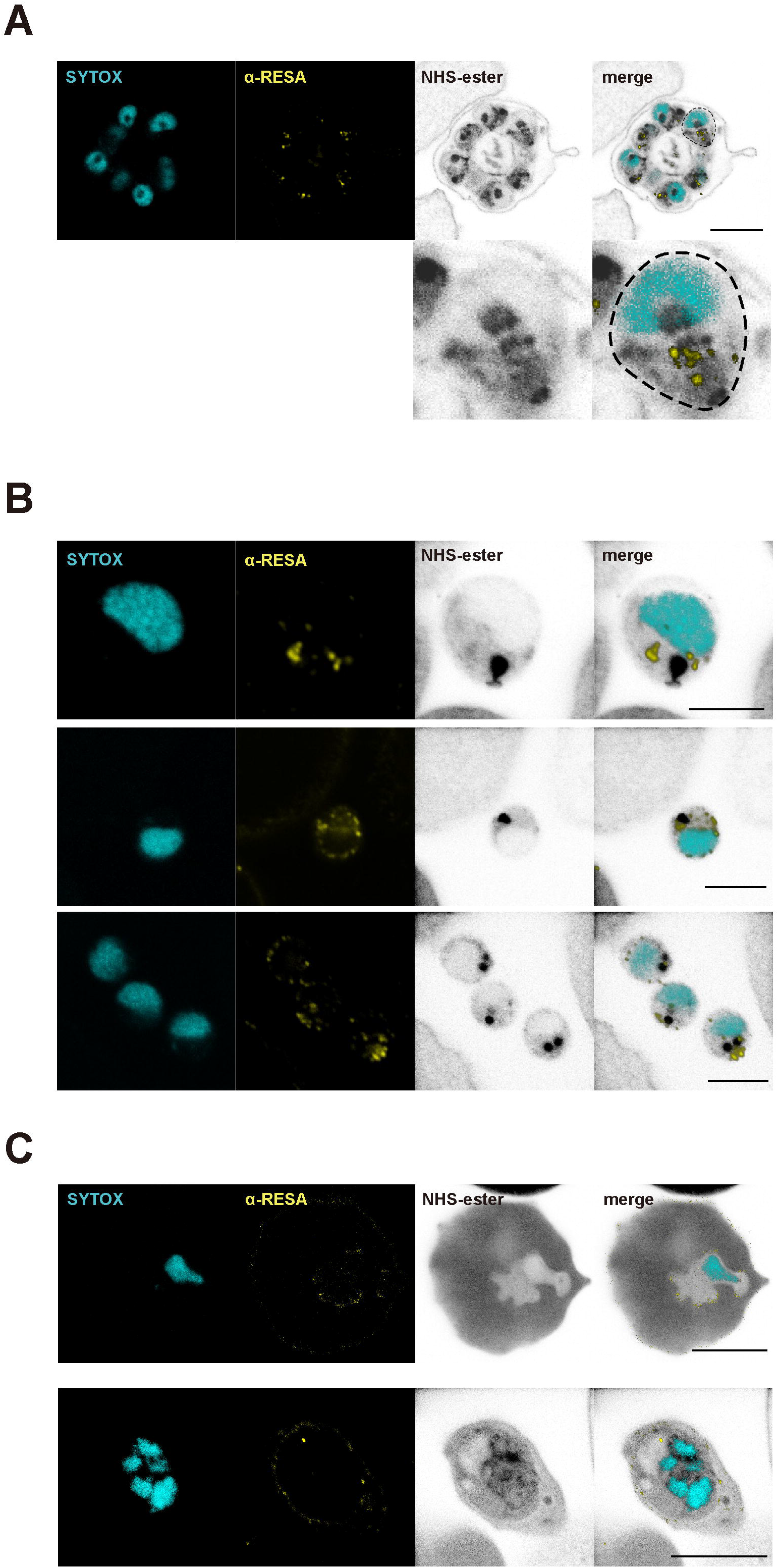
Anti-RESA antibody described the detailed locations of dense granules throughout the asexual life cycle of parasites. Representative U-ExM images of segmented schizonts (A), merozoites (B), rings (C, upper panel), and early schizonts (C, lower panel) labeled with anti-RESA antibody. Protein density and nuclei were visualized with NHS ester Alexa Fluor 405 and SYTOX Deep Red, respectively. RESA, amines, and nuclei are shown in yellow, grayscale, and cyan, respectively. Scale bars = 10 μm (A,C) and 5 μm (B).

In conclusion, this study used U-ExM to detect RESA in protein-condensed small globular organelles which were likely DG, in accordance with previous IEM observations. U-ExM is compatible with conventional confocal laser-scan microscopy systems. Therefore, U-ExM will significantly improve the identification of novel DG proteins, and make the process more efficient than traditional IEM methods; thus, aiding in describing parasitism in erythrocytes by *P. falciparum*.

## Acknowledgements

We appreciate the Japanese Red Cross Society for providing human erythrocytes. This work was supported in part by JSPS KAKENHI (Grant numbers JP 21K15429, JP 21KK0138, and JP 23KK0140). The funders had no role in the study design, data collection and analysis, decision to publish, or preparation of the manuscript. We thank Thomas J. Templeton for his critical reading of this manuscript.

## Author contributions

Conceptualization: J.F., T.T., and E.T. Data collection: J.F. and E.T. Funding acquisition: J.F., T.T., and E.T. Writing of original draft: J.F. and E.T. Writing review and editing: J.F., T.T., and E.T.

