## Supplementary Figure for "Ultrastructural expansion microscopy (U-ExM) visualization of malaria parasite dense granules using RESA as a representative marker protein"

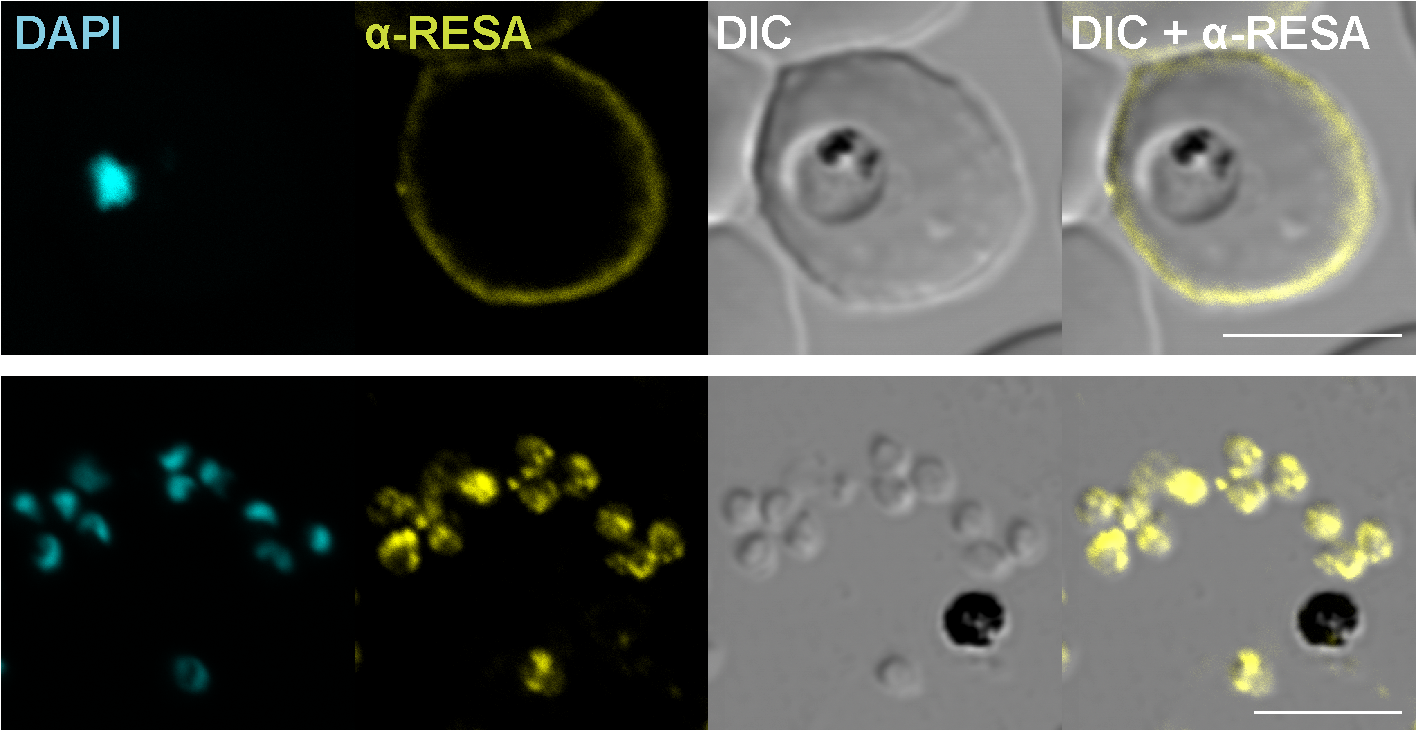

**Supplementary Figure 1. RESA antibody reacted with RESA in the malaria parasite using conventional immunofluorescence assay.**

Representative immunofluorescence assay images of ring and trophozoite stage parasites labeled with anti-RESA antibody. Nuclei were stained with DAPI. RESA and nuclei are shown in yellow and cyan, respectively. Scale bars = 5 μm.
