## Supplementary Methods for "Ultrastructural expansion microscopy (U-ExM) visualization of malaria parasite dense granules using RESA as a representative marker protein"

**Parasite culture**

Plasmodium falciparum 3D7 strain parasites were cultured in O^+^ erythrocytes at 2% hematocrit in RPMI-1640 media with L-glutamine (Thermo Fisher Scientific, Waltham, MA, USA) supplemented with 5.94 mg/mL HEPES (Sigma–Aldrich, Burlington, MA, USA), 0.05 mg/mL hypoxanthine (Sigma–Aldrich), 0.225% sodium bicarbonate (Thermo Fisher Scientific), 10 μg/mL gentamicin (Thermo Fisher Scientific), and 0.5% Albumax II (Thermo Fisher Scientific) and maintained at 37°C in a gas mixture of 5% O_2_ , 5% CO_2_, and 90% N_2_. Human erythrocytes were obtained from the Japan Red Cross Society (Tokyo, Japan) (#25J0176). The experiments with human erythrocytes were conducted under the guidelines of the ethics committee of Ehime University (Aidaiibyourin#1712001).

**Mouse anti-RESA monoclonal antibody**

Hybridoma clones producing mouse anti-RESA monoclonal antibodies (mAbs) were generated and purchased from Kitayama Labes (Ina, Japan) as described [1]. Briefly, female BALB/c mice 8 weeks old were immunized in their foot pads twice with 50 μg of synthetic RESA peptide (EENVEENVEENVEENVEENV) conjugated with keyhole limpet hemocyanin (KLH) formulated with Freund’s complete adjuvant for the priming and incomplete adjuvant for the boost two weeks apart. A final intravenous boost with the unconjugated RESA peptide in PBS was administered six weeks after the boost. Lymphocytes from the inguinal lymph nodes were used to fuse with P3-X63-Ag8-U1 myeloma cells to produce hybridoma cells. Culture supernatants from hybridomas were screened for reactivity against unconjugated RESA peptide. Positive hybridoma cells were cloned, and the antibody isotypes were determined as described [1]. Cloned cell lines were expanded in a serum-free culture medium, and immunoglobulin G (IgG) was purified using an MAbTrap kit (GE Healthcare, Camarillo, CA). We confirmed the reactivity and specificity of purified mAb in an indirect immunofluorescence assay (IFA), in which the immunofluorescence stain signal was consistent with previous studies (Supplementary Figure 1).

**Immunofluorescence assay**

Infected RBCs were immobilized on 22 x 22 mm coverslips (thickness No. 1.5H) (Thorlabs, Inc., Newton, NJ, USA) coated with poly-D-lysine (Sigma–Aldrich). After the immobilization, samples were fixed with 4% paraformaldehyde in phosphate buffered saline (PBS) for 10 min and 50 mM dimethyl suberimidate dihydrochloride (Sigma–Aldrich) in borate buffer for 20 min, or 4% paraformaldehyde/0.075% glutaraldehyde in PBS followed by quenching with 10 mM glycine in PBS for 30 min. Fixed samples were permeabilized with 0.3% Triton X-100 in PBS and blocked with 1% skim milk in 0.05% Tween 20 in PBS for 60 min. Samples were treated with mouse anti-RESA mAb (4F12F3) (1:2000 dilution) overnight and Alexa Fluor 488-conjugated goat anti-mouse (1: 1000 dilution) together with DAPI for 60 min, followed by sealing with Antifade Mounting Medium. Samples were observed with a 60× objective using an LSM710 confocal laser scanning microscope (Carl Zeiss, Oberkochen, Germany).

**Ultrastructural expansion microscopy (U-ExM)**

Infected RBCs were immobilized on 12 mm round coverslips (MATSUNAMI, Osaka, Japan) coated with poly-D-lysine (Sigma–Aldrich) in 12-well plates. After the immobilization, samples were fixed with 1 mL of 4% paraformaldehyde in PBS for 30 min. After the fixation, coverslips were washed three times with pre-warmed (37°C) PBS. For protein anchoring, coverslips were treated with 1 mL of 1.4 % formaldehyde/2% acrylamide (FA/AA) in PBS at 37°C overnight. After removal of FA/AA and one wash with PBS, coverslips were put on activated monomer solution (19% sodium acrylate (BLDpharm), 10% acrylamide (Sigma–Aldrich), 0.1% N,N’-methylenebisacrylamide (Sigma–Aldrich), 0.25% N,N,N',N'-Tetramethylethylenediamine (TEMED) (Thermo), and 0.25% ammonium persulfate (Thermo)) facing the side that the cells were adhere to and gelated at 37°C for 30 min. After gelation, gels on coverslips were put in 6-well plates containing denaturation buffer (200 mM sodium dodecyl sulfate (SDS), 200 mM NaCl, 50 mM Tris-base, pH 9) and incubated for 15 min with shaking. Separated gels were transferred into 1.5 mL tubes containing denaturation buffer and boiled at 95°C for 90 min to ensure maximal expansion. For the 1st expansion, boiled gels were put into 10 cm Petri dishes containing 25 mL of MilliQ water and shook for 30 min three times. The expanded gels were shrunk with 25 mL of PBS and shook for 15 min two times. Shrunk gels were transferred into 6-well plates containing 3% BSA in PBS and blocked for 30 min at room temperature. After blocking, the gels were incubated with mouse anti-RESA (4F12F3), or rabbit anti-RAP1 or anti-AMA1 polyclonal antibodies [2] (1:100 dilution) in 3% BSA in PBS overnight. Gels were washed three times with 0.5% Tween 20 in PBS for 10 min each time, followed by an incubation with Alexa Fluor 488-conjugated goat anti-mouse or anti-rabbit (1:500 dilution) together with 8 µM NHS ester Alexa Fluor 405 and 1 µM SYTOX Deep Red in PBS for 2.5 hours. Following three washes for 10 min with 0.5% Tween 20 in PBS, the gels were transferred into 10 cm Petri dishes for re-expansion with three 30 min washes of MilliQ water. Re-expanded gels were cut into roughly 10 mm x 10 mm sections and mounted on a poly-D-lysine (Sigma–Aldrich) coated glass bottom dish for imaging. Samples were observed with a 60× objective using an LSM710 confocal laser scanning microscope (Carl Zeiss MicroImaging,

Thornwood, NY, US).
